# Shoc2 controls ERK1/2-driven neural crest development by balancing components of the extracellular matrix

**DOI:** 10.1101/2022.06.13.495941

**Authors:** Rebecca G. Norcross, Lina Abdelmoti, Eric C. Rouchka, Kalina Andreeva, Olivia Tussey, Daileen Landestoy, Emilia Galperin

## Abstract

The extracellular signal-regulated kinase (ERK1/2) pathway is essential in embryonic development. The scaffold protein Shoc2 is a critical modulator of ERK1/2 signals, and mutations in the *shoc2* gene lead to the human developmental disease known as Noonan-like syndrome with loose anagen hair (NSLH). The loss of Shoc2 and the *shoc2* NSLH-causing mutations affect the tissues of neural crest (NC) origin. In this study, we utilized the zebrafish model to dissect the role of Shoc2-ERK1/2 signals in the development of NC. These studies established that the loss of Shoc2 significantly altered the expression of transcription factors regulating the specification and differentiation of NC cells. Using comparative transcriptome analysis of NC-derived cells from *shoc2* CRISPR/Cas9 mutant larvae, we found that Shoc2-mediated signals regulate gene programs at several levels, including expression of genes coding for the proteins of extracellular matrix (ECM) and ECM regulators. Together, our results demonstrate that Shoc2 is an essential regulator of NC development. This study also indicates that disbalance in the turnover of the ECM may lead to the abnormalities found in NSLH patients.

## INTRODUCTION

The Ras-ERK1/2 canonical signaling pathway is activated by multiple extracellular cues and promotes cell proliferation, cell cycle progression, and a myriad of other cellular functions. The pleiotropic role of the ERK1/2 signals in various tissues has been well described by a large body of literature. Mutations in genes of the Ras-ERK1/2 signaling pathway can result in congenital disorders that are collectively termed RASopathies (Rauen, 2013, Tidyman and Rauen, 2009). Individual RASopathies display considerable variability within their clinical phenotypes. The hallmarks of RASopathies include distinct facial features, cardiac defects, growth delays, neurologic issues, and gastrointestinal difficulties (Tajan et al., 2018, Riller and Rieux-Laucat, 2021). Given the universal significance of the ERK1/2 pathway in all cell types, it is somewhat surprising that the clinical hallmarks of RASopathies are very specific. Thus, to develop therapeutic avenues for minimizing these diseases, it is essential to understand the precise etiology and pathogenesis of individual malformation syndromes. This requires an in-depth understanding of the functional developmental roles played by genes that are mutated in specific disorders.

Noonan-like syndrome with loose anagen hair (NSLH; OMIM: 607721) is a relatively rare RASopathy with an estimated incidence of 1:50000 live births (Mazzanti et al., 2003). NSLH is characterized by cardiac (most often valvuloseptal) abnormalities, proportional short stature, facial dysmorphia (e.g. ocular hypertelorism), and a variety of less penetrant defects (e.g. cognitive, genitourinary, auditory abnormalities, and unique loss of anagen hair). The majority of NSLH patients carry autosomal dominant mutations in the *shoc2* gene that encodes the non-enzymatic leucine-rich repeat scaffold protein Shoc2 (Cordeddu et al., 2009). Shoc2 plays a key role in the amplification of the signals of the ERK1/2 pathway and its loss is detrimental to embryonic development (Jang et al., 2020).

Studies from several labs demonstrated that to modify ERK1/2 signals Shoc2 assembles an intricate protein machinery (Jang and Galperin, 2016). This protein machinery facilities the amplification of intracellular signals transmitted *via* the Shoc2 module. It also provides a negative feedback mechanism to fine-tune the transient Shoc2-mediated signals (Jang et al., 2014, Jang et al., 2019a, Wilson et al., 2021, Jang et al., 2015). Yet, the question of what biological activities are regulated by the Shoc2-transmitted signals remains elusive. The ablation of Shoc2 in mice leads to early embryonic lethality and partial embryo absorption at E8.5 (Yi et al., 2010). In our earlier studies, we employed a powerful zebrafish vertebrate model to generate a heritable CRISPR/Cas9 knock-out of Shoc2 (Jang et al., 2019b). The loss of Shoc2 in zebrafish induced an array of developmental defects. Interestingly, the most prominent deficiencies in morphogenesis of the Shoc2 *null* mutants were in the tissues of neural crest (NC) origin: facial cartilage, bone, and pigment (Jang et al., 2019b). Importantly, these deficiencies phenocopied the cranio-skeletal anomalies observed in humans with NSLH (Jang et al., 2019b). Moreover, anomalies of patients with Shoc2-related NSLH are well-aligned with symptoms of neurocristopathies - pathologies resulting from the abnormal specification, migration, differentiation, or death of NC cells (NCCs) during embryonic development (Vega-Lopez et al., 2018).

NCCs are stem cell-like populations unique in their capacity to migrate a great distance during embryonic development (Martik and Bronner, 2021). These cells give rise to different types of critical tissues, including craniofacial cartilage and bone, heart muscle, pigment, peripheral neurons, and ganglia (Mendez-Maldonado et al., 2020). The events of NC formation and differentiation into a variety of derivatives are tightly controlled by the activation of the early gene regulatory network (Martik and Bronner, 2017). ERK1/2 pathway signals are essential for the establishment of NCC populations (Geary and LaBonne, 2018, Newbern et al., 2008, Dinsmore and Soriano, 2018) and are necessary to control the timing of the progression from pluripotency to lineage restriction (Parada et al., 2015, Newbern et al., 2011). The defects related to NCC migration and specification have also been suggested to underlie changes in anatomical structures of mice models of RASopathies such as Noonan syndrome (Nakamura et al., 2009a, Stewart et al., 2010, Lajiness et al., 2014) and Neurofibromatosis 1 (Nakamura et al., 2009b, Grossmann et al., 2009, Padmanabhan et al., 2009). Together, these and other studies indicated that NCCs exhibit a specific threshold sensitivity to the deficiencies in ERK1/2 signals during morphogenesis. Yet, a clear mechanistic link between any RASopathy-causing mutations and the resulting developmental defect is still missing.

Here, we leverage the power of the zebrafish model to delineate the role of Shoc2 in the development of the NC. We establish that Shoc2 deficiency affects early NC gene expression and demonstrate that Shoc2-guided signals are critical for the cell fate determination of NC derivatives. The loss of Shoc2 affects NC-derived precursors as well as differentiated populations of craniofacial cartilage and cranial ganglia. We found that Shoc2 is critical for the differentiation of pigment cells and myelinated Schwann cells. Moreover, we observed that transcriptional circuits of *sox10*-positive cells derived from CRISPR/Cas9 Shoc2 *null* mutants were markedly altered. The loss of Shoc2 leads to perturbations in the expression of the extracellular matrix (ECM) components and proteins associated with ECM. These perturbations are likely to be responsible for the cranio-skeletal defects observed in the Shoc2 CRISPR/Cas9 *null* mutants. Most significantly, our results point to a role for Shoc2 as a novel regulator of NC and early embryonic cell fate programming.

## RESULTS

### Loss of Shoc2 alters neural plate border gene expression

Our earlier studies characterized the overall developmental abnormalities of two Shoc2 mutant zebrafish lines generated by CRISPR/Cas9 mutagenesis: *shoc2*Δ*22* and *shoc2*Δ*14* (Jang et al., 2019b). These studies demonstrated that the developmental abnormalities found in CRISPR/Cas9 *shoc2* mutants were in several neural crest (NC) derived tissues, including malformations of multiple cartilage elements, bone, and pigment (**Fig. S1A**). Here we investigate the effect of Shoc2-mediated signals on the development of NC further. Homozygous *shoc2* CRISPR/Cas9 mutant embryos are phenotypically distinguishable from their control siblings only at 5 days post fertilization (dpf), likely due to the contribution of the maternal *shoc2* mRNA. Thus, we utilized a targeted morpholino oligonucleotide (MO)-mediated knock-down for an acute depletion of Shoc2. The efficacy of the *shoc2* MO to interfere with the translation of *shoc2* mRNA was validated by Western blot analysis using an anti-Shoc2 antibody. The results of these experiments were analogous to what we reported earlier (Wilson et al., 2021) (**Fig. S1B**). The specificity of the *shoc2* MO was also validated in our earlier studies, when co-injection of wild-type (WT) human *shoc2* mRNA with *shoc2* MO partially rescued *shoc2* MO-induced deficits (Jang et al., 2019b, Wilson et al., 2021).

First, we analyzed whether the loss of Shoc2 affects the expression of genes involved in the definition of the neural plate border (NPB) territory that gives rise to the NC precursors (Rocha et al., 2020, Stuhlmiller and Garcia-Castro, 2012). The expression of transcription factors specific to the NPB, PR domain containing 1a, with the ZNF domain *(prdm1a)* and paired box 7 *(pax7)* (Olesnicky et al., 2010, Hernandez-Lagunas et al., 2005, Roellig et al., 2017, Powell et al., 2013, Vadasz et al., 2013), was examined at the 2-somite stage, when NC progenitors are present in the anterior portion of the embryos, using whole body *in situ* hybridization (WISH). In the *shoc2* morphant embryos, the *prdm1a* expression pattern was altered and the lateral edge of the neural plate was farther apart from the adaxial cells adjacent to the midline compared to control larvae (**Fig. 1A, E**). The *shoc2* morphants also had a mild alteration in the expression pattern of *pax7* (**Fig. 1B, E**). Furthermore, cells expressing the forkhead-box transcription factor *foxd3,* an essential “NC specifier”, were spaced farther from the midline in the *shoc2* morphants than in control embryos. The expression of *foxd3* was also reduced in the posterior region (**Fig. 1C, E**). Although, the loss of Shoc2 mildly effected on the expression pattern of the pan-neural marker SRY-box transcription factor 2 (*sox2),* the *sox2* domain was wider in the *shoc2* morphants than in control larvae (**Fig. 1D, E**). We have not detected changes in the major-to-minor axis ratio of the *shoc2* morphants at 11 hours post fertilization (hpf), suggesting that convergent extension cell movement during gastrulation was largely unaffected. Thus, it is possible that the expression of Shoc2 is required in the very early steps of NC development- the definition of the NPB territory.

**Figure 1.**
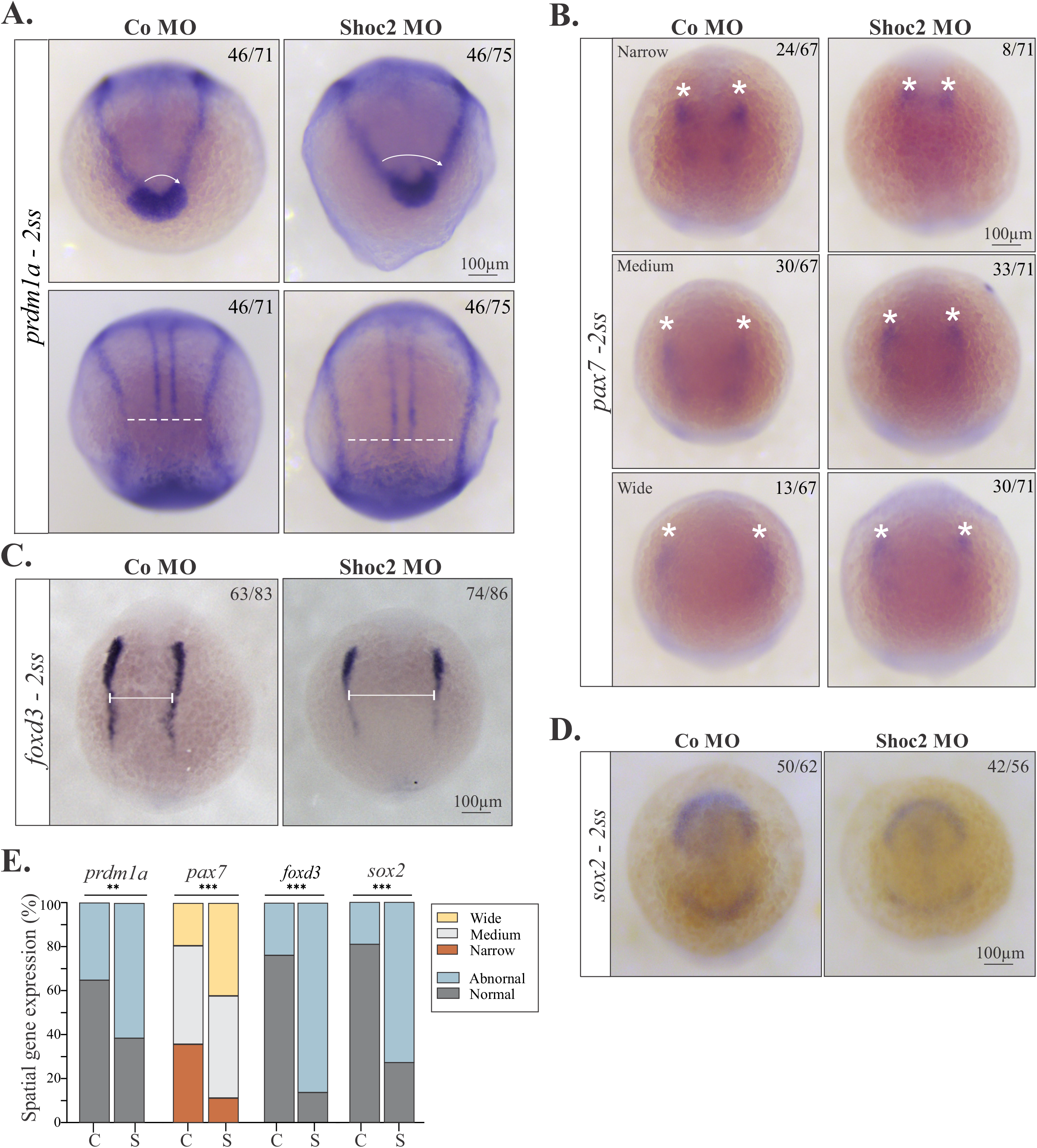
Analysis of gene expression at NPB of *shoc2* morphants. Dorsal views of control and *shoc2* morphant embryos showing expression of *prdm1a* (**A**), *pax7* (**B**), *foxd3* (**C**), and *sox2* (**D**) in 2-somite-stage embryos. Aberrant expression patterns of *prdm1a*, *pax7*, *foxd3*, and *sox2* are evident in *shoc2* morphants. Arrows, lines, and asterisks indicate parameters that were assessed to determine the abnormal expression patterns. The graph (E) shows the frequency of observed patterns from at least three independent experiments. The total number of embryos is indicated on each image. Statistically significant differences between *shoc2* MO and control MO according to the Pearson’s chi-squared test are indicated by *p<0.05, **p<0.01, ***p<0.001

### Shoc2 is necessary for induction of neural crest

To understand whether the loss of Shoc2 affects the specification of cells in the NPB to the NC fate, we used WISH to analyze the expression of the “neural crest specifiers” *foxd3*, snail family zinc finger 2 *(snai2)*, and SRY-box transcription factors *sox10*, and *sox9a* at 5 somites, before the onset of NC migration (Stuhlmiller and Garcia-Castro, 2012, Li and Cornell, 2007). We found that cells expressing *foxd3* (**Fig. 2A, F**) and *sox10* (**Fig. 2B, F**) were shifted more laterally from the midline in embryos injected with Shoc2 MO. The expression levels of *sox10* appeared to be reduced (**Fig. 2B, F**). The loss of Shoc2 also affected the expression patterns of *snai2* and *sox9a* (**Fig. 2C, D and F**). Moreover, we found a marked reduction in expression of *crestin,* a pan-NC marker (Luo et al., 2001, Rubinstein et al., 2000), in the anterior portion of NPB of Shoc2 morphants (**Fig. 2E, F**). These findings suggest that the NC is specified in the *shoc2* morphants, but is not patterned correctly.

**Figure 2.**
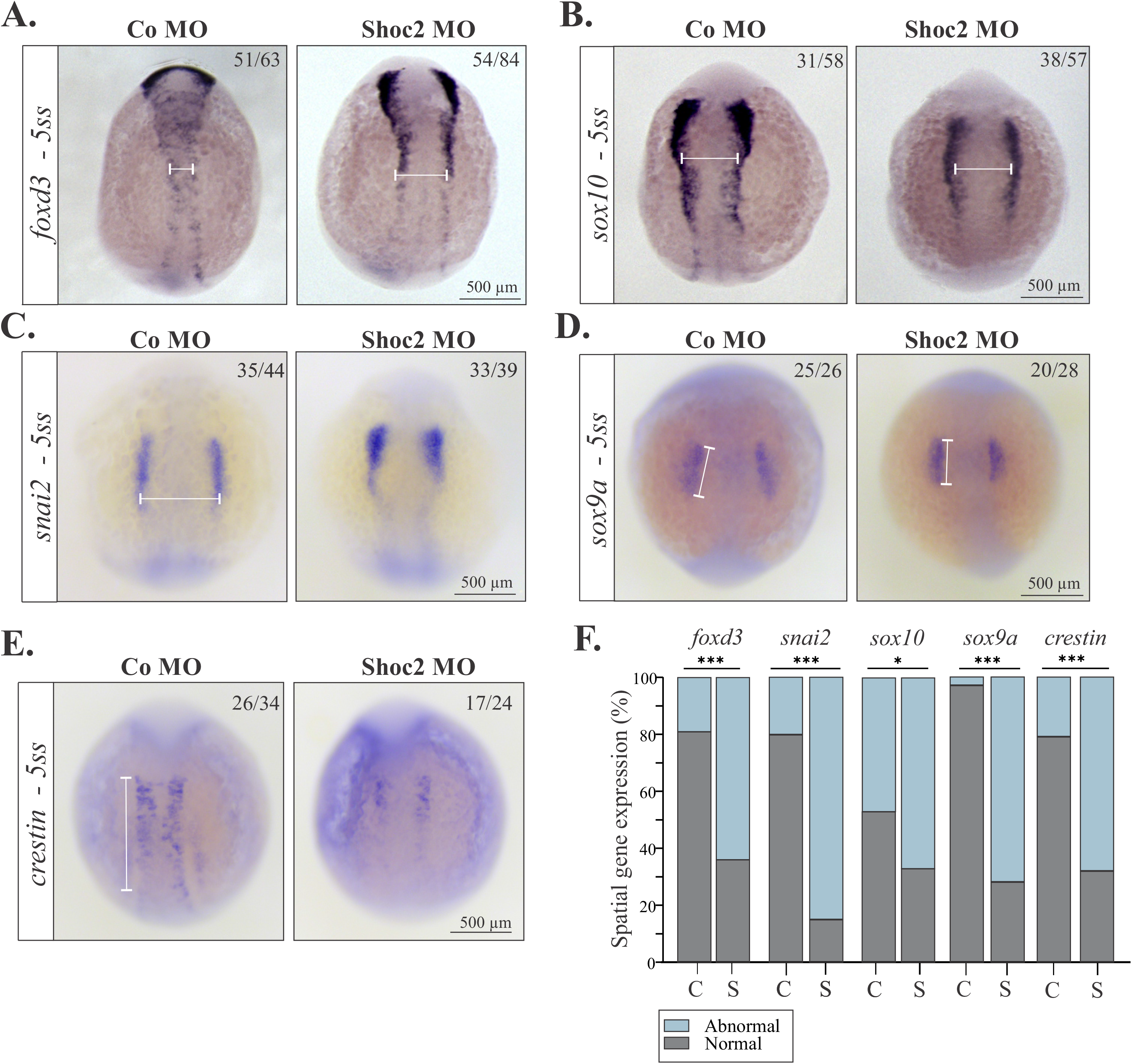
Molecular defects in NC specification in *shoc2* morphants. Dorsal views of control and *shoc2* morphant embryos show expression of NC specifiers *foxd3* (**A**), *sox10* (**B**), *snai2* (**C**), *sox9a* (**D**) and *crestin* (**E**) in 5-somite-stage embryos. White lines indicate the parameters that were assessed to determine the changes in expression of *foxd3*, *sox10*, *snai1*, *sox9a* and *crestin*. The graph (**F**) shows the frequency of observed expression patterns (C) from at least three independent experiments. The total number of embryos used in the statistical analysis is indicated on each image. Statistically significant differences between *shoc2* MO and control MO according to the Pearson’s chi-squared test are indicated by *p<0.05, **p<0.01, ***p<0.001.

### Shoc2 effect on the migratory neural crest cells

Following NC induction and specification, the NCCs transition to become actively migrating mesenchymal cells (Rocha et al., 2020). This gradual transition requires the regulation of many effector genes and relies on transcription factors that are also involved in NCC fate specification, such as *foxd3* and *snai2* (Stewart et al., 2006, Barrallo-Gimeno and Nieto, 2005). Thus, we assessed whether expression of *foxd3* and *snai2* was affected by the loss of Shoc2 at the 18-somite stage (ss). In *shoc2* morphant larvae, the spatiotemporal migratory paths of cells expressing *foxd3* and *snai2* were altered in both the cranial and trunk NC of 18-somite *shoc2* morphants (**Fig. 3A, C)**. The expression of *foxd3* was decreased in somites, in the cranial NC around the otic vesicle and the late migrating NC precursors. We also detected reduced expression of *snai2* in migrating NC precursors. Consistent with reduced expression of *foxd3* and *snai2* in migratory NCCs, a significant loss in NCCs expressing *crestin* was detected in the first trunk segments of the migrating ventral streams between the neural keel and somites of *shoc2* morphants (**Fig. 3A, C (bottom panels).** Together, our findings point to either a loss in migratory NCCs or to defects in the premigratory NCC maintenance of the *shoc2* morphants.

**Figure 3.**
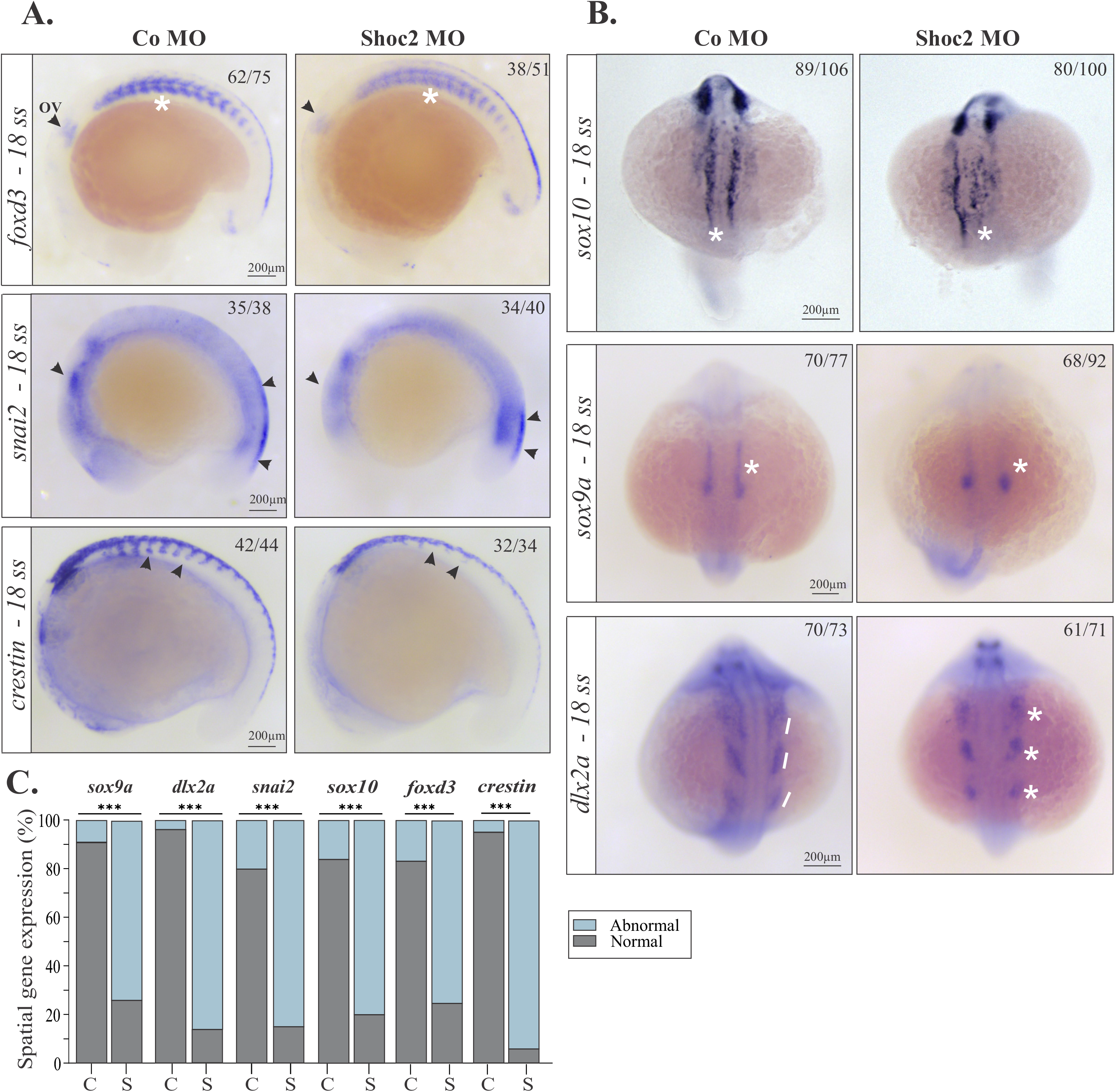
Gene expression abnormalities in early migrating NCCs in *shoc2* morphants. Lateral views of control and *shoc2* morphant embryos showing expression of (**A**) *foxd3, snai2,* and *crestin* at the 18-somite stage. (**B**) Dorsal views of control and *shoc2* morphant embryos showing expression of *sox10, sox9a,* and *dlx2a*. Arrows, asterisks, and lines indicate the parameters that were assessed to determine the abnormal expression patterns. The graph (**C**) shows the frequency of observed abnormal patterns after injection of *shoc2* MO or a control MO from at least three independent experiments. The total number of embryos assessed is indicated in each image. Statistically significant differences between *shoc2* MO and control MO according to the Pearson’s chi-squared test are indicated by *p<0.05, **p<0.01, ***p<0.001. ov: otic vesicle.

To better understand the extent of Shoc2 loss of function on the specification of non-ectomesenchymal NC derivatives and their migration, the expression of the transcription factors *sox10 and sox9a* was assessed (Yan et al., 2005). Compared to control larvae, *shoc2* morphants exhibited irregular distribution of the anterior rostal-to-caudal *sox10*-positive NCCs with a greatly diminished area of cells positive for the expression *sox9a* (**Fig. 3B, C**). The 18 ss of zebrafish development generally correlates with initial migration of cranial NCCs. Examination of the expression of transcription factor distal-less homeo-box 2 (*dlx2a)* demonstrated that in *shoc2* morphants three streams of mature cranial NCCs migrating towards the prospective pharyngeal arches (Sperber et al., 2008) were noticeably diminished (**Fig. 3B, C (bottom panels**).

Mesenchymal transition of NCCs also involves cell surface changes, dissolution of cadherin-mediated adherens junctions and a “cadherin switch” from E-cadherin to N-cadherin (Theveneau and Mayor, 2012). While the pre-migratory NCCs in zebrafish mostly express E-cadherin (*cdh1*), migratory NCCs predominantly express N-cadherin (*cdh2*) (Taneyhill and Schiffmacher, 2017, Scarpa et al., 2015). The impact of the Shoc2 loss on the migratory properties of the NCCs was determined by analyzing the expression of *cdh1*, *cdh2* and additional genes expressed in NCCs at the time of epithelial-to mesenchymal transition, *twist1a* and *snai2,* using qRT-PCR. Data in **Fig. S2A** establish that at 24 hpf the expression of *chd1*, *cdh2*, and *twist* were significantly reduced in *shoc2* morphants. This suggests that while *shoc2* function is necessary to control expression of *chd1*, *cdh2,* and *twist1a*, it does not seem to be necessary to control a “cadherin switch”. Importantly, differences in expression of genes regulating migration of NCCs also coincided with a dramatic decrease in migrating *crestin*- and *foxd3*-positive cells (**Fig. S2B-D and Fig 3A)**. Together, our data suggest that loss of Shoc2 leads to defective NCC specification and migration, which, in turn, could underlie other developmental defects observed in zebrafish *shoc2* mutants.

### Shoc2 signals in the development of pigment cells

Zebrafish NCCs are fate-restricted from an early stage, and individually labeled premigratory NCCs typically produce differentiated cells of only a single class (Rocha et al., 2020). NCCs differentiate into the cells of the peripheral nervous system, the craniofacial osteocytes and chondrocytes, and three major pigment cells types: xanthophores, iridophores, and melanocytes (Rocha et al., 2020). Our earlier studies reported that ablation of Shoc2 affected the pigmentation pattern of Shoc2 *null* larvae and, at 6 dpf, *shoc2* mutants lost the regularity of melanophore patterning and presented with overlapping lentigines (Jang et al., 2019b). Here, we demonstrate additional deficiencies in the development of pigment cells resulting from the loss of Shoc2. Compared to WT larvae, numbers of iridophores in Shoc2 *nulls* were reduced substantially, particularly in the trunk and tail, at 120 hpf (**Fig. S3)**, indicating defects in the differentiation of pigment NC lineage.

### Shoc2 and the peripheral nervous system

Neurogenic derivatives of NCCs are generated at all axial levels (Rocha et al., 2020). In the cranial region, NCCs contribute to the cranial ganglia, Schwann, and satellite cells, while in the trunk they give rise to sensory neurons of the dorsal root ganglia (DRG), sympathetic neurons, and Schwann cells (Etchevers et al., 2019). To determine the extent of Shoc2 requirement for the development of the peripheral nervous system, we evaluated the expression of the transcription factor *foxd3* at 48 hpf when it is required for the proper specification of the peripheral neurons of DRG, enteric neuron, and the cranial ganglia precursors (Wang et al., 2011) (**Fig. 4A, E**). In control larvae, *foxd3* expression was easily detected in the cranial ganglia-associated glia of the developing trigeminal ganglion, the pre- and post-otic ganglia, and the trunk satellite glia associated with DRG. However, in the embryos injected with *shoc2* MO, the expression of *foxd3* was greatly diminished in the ganglia-associated glia (**Fig. 4A, E**). The *shoc2* morphants also had irregular *foxd3*-positive ventral streams of NCCs and the DRG structures, including abnormal ectopic expression patterns of precursors of the peripheral nervous system (**Fig. 4A,** inset).

**Figure 4.**
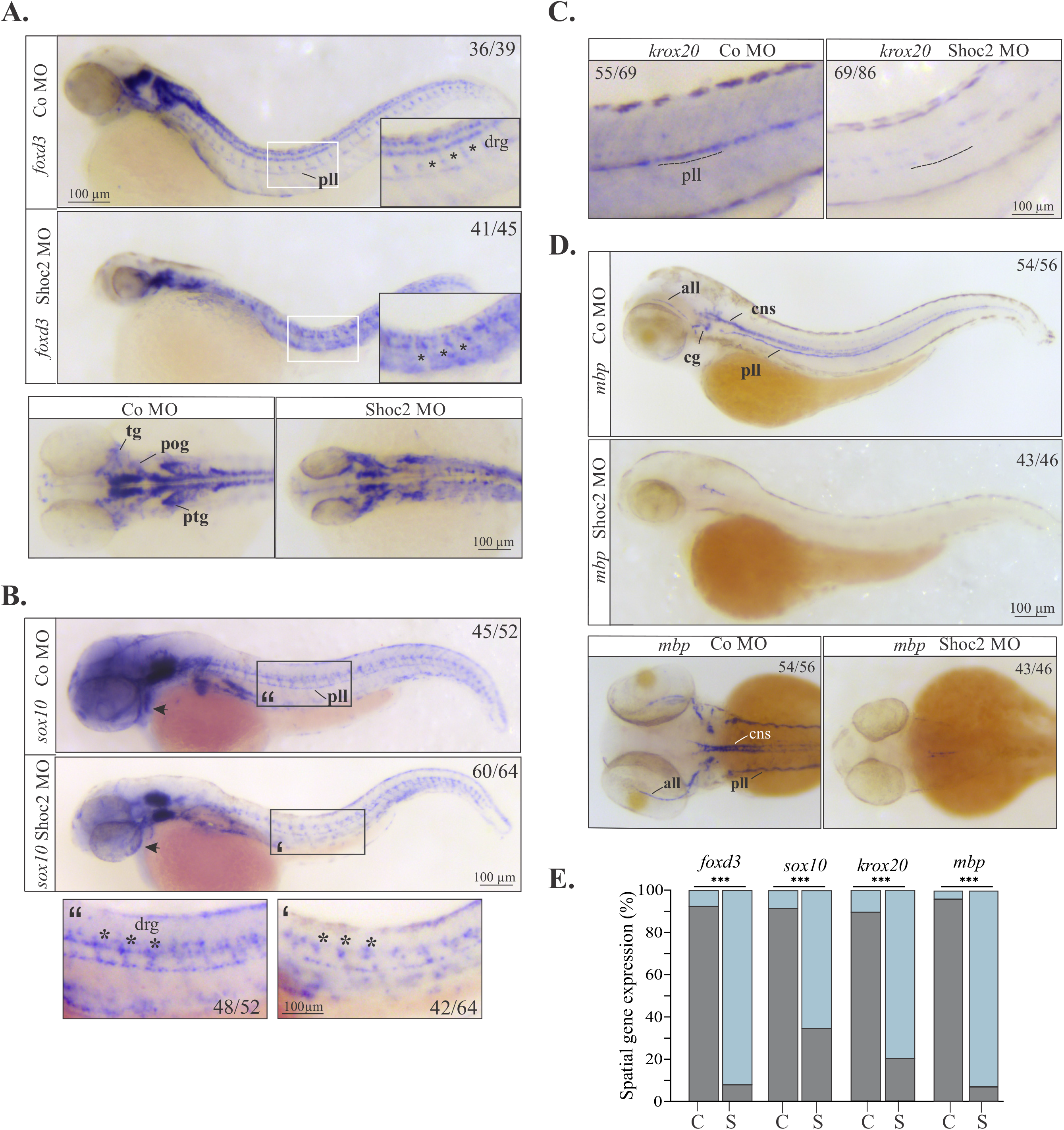
Molecular defects in NC specification and differentiation in *shoc2* morphants. Lateral and dorsal views of control and *shoc2* morphant embryos show the expression of *foxd3* (**A**) and *sox10* (**B**) at 2dpf, and *krox20* (**C**) and *mbp* (**D**) at 3dpf. Asterisks and lines indicate the parameters that were assessed to determine the abnormal expression patterns. The graph (**E**) shows the frequency of abnormal patterns from at least three independent experiments. The total number of embryos assessed is indicated in each image. Statistically significant differences between *shoc2* MO and control MO according to the Pearson’s chi-squared test are indicated by *p<0.05, **p<0.01, ***p<0.001. all: anterior lateral line. cns: central nervous system. pll: posterior lateral line. cg:cranial ganglia. drg: dorsal root ganglia. pog: preotic ganglia; ptg: postotic ganglia; tg: trigeminal ganglia

To assess further whether Shoc2-mediated signals are required for the late-forming neural progenitors (neurons and glia) in the trunk, we examined the expression of the *sox10* gene at 48 hpf (**Fig. 4B, E**). The loss of Shoc2 resulted in a dramatic reduction of glial cells and NC-derived sensory neurons of DRGs in zebrafish embryos. We found that, similarly to *foxd3*, *sox10* expression was lost in segmentally arranged lines of cells laying adjacent to the notochord (the precursors of the peripheral nervous system) in the trunk of the *shoc2* morphants (**Fig. 4B (**inset**), E**). The loss in expression of *foxd3* and *sox10* in *shoc2* morphant larvae indicates that Shoc2 contributes to the specification of the NC peripheral neurogenic progenitor population.

To understand better the role of Shoc2 in the development of the peripheral nervous system, we assessed a subset of the glia, the Schwann cells, that surround the ganglia of the lateral line and ensheath the lateral line nerves (Brosamle and Halpern, 2002). We examined the expression of transcription factor *krox20,* which, together with *sox10*, regulates terminal differentiation by controlling expression of another marker of glial differentiation, *myelin basic protein (mbp).* Compared to controls (**Fig. 4C, E**), the *shoc2* morphants showed a decrease in overall *krox20* expression which was particularly clear along the lateral line (**Fig. 4C**). Likewise, we found a dramatic reduction in expression of *mbp* in the anterior (all) and posterior lateral line (pll), cranial ganglia and central nervous system of *shoc2* morphants at 3 dpf (**Fig. 4D, E**).

To determine if overall reduction in the differentiation of NCCs into glial and pigment derivatives was due to increased apoptosis, TUNEL labelling was carried out. In contrast to control larvae, which exhibited limited staining, embryos injected with the *shoc2* MO showed an increase in apoptosis along the trunk (at 24 hpf) and in the hindbrain region (at 48 hpf) (**Fig. S4A-C**). We also assessed cell proliferation by labeling larvae with anti-phospho-Histone H3 antibody and Alexa Fluor 488 dye at 24 hpf. We have not detected changes in numbers of GFP-positive cells, indicating that Shoc2 signals do not affect cell proliferation (**Fig. S4D, E**). These data suggest that increased apoptosis may partially account for the reduction of *foxd3* and *sox10* expression in specified NCCs cells. The increase in apoptotic cells in *shoc2* morphants also suggest that Shoc2 signals contribute to the survival of certain cell populations.

### Shoc2 regulates the expression of cranial NCC (cNCC) specific genes in the posterior pharyngeal arches

One of the strongest abnormalities of the Shoc2 CRISPR/Cas9 mutants is the defects in the cartilaginous structures of the viscerocranium (**Fig. S1A**) (Jang et al., 2019b). Data in **Fig. 3B** strongly indicate that *dlx2a*-positive NCCs fated to become CNCCs do not migrate properly in *shoc2* morphant embryos, possibly affecting formation of the cartilaginous structures of the viscerocranium. Moreover, we also found that chondrocyte stacking within the cartilaginous elements of *shoc2* morphants was not as orderly arrayed as in controls (**Fig. 5A**). Thus, to better understand the craniofacial phenotypes of Shoc2 mutant larvae, we examined the differentiation of post-migratory CNCCs into chondrocytes. WISH was used to analyze the expression of the NC and prechondrogenic marker *sox9a* critical for cartilage morphogenesis and chondrocyte stacking (Yan et al., 2005). Compared to embryos injected with the control MO, in which the *sox9a* expression was readily detectable in cranial structures and in fin buds, *sox9* expression was decreased in various cranial structures of *shoc2* morphants and was mostly absent from the pectoral fin (**Fig. 5B**). In contract to the decreased *sox9a* expression in cranial structures, *sox9a* expression was elevated in the trunk of the Shoc2 morphant embryo. These data, validated by the qPCR analysis (**Fig. 5C**), suggest that loss of Shoc2 has region-specific effects on *sox9* gene expression, and that the cartilage abnormalities of *shoc2* mutants are likely due to the defective specification and migration of cranial NCCs or chondrocytes.

**Figure 5.**
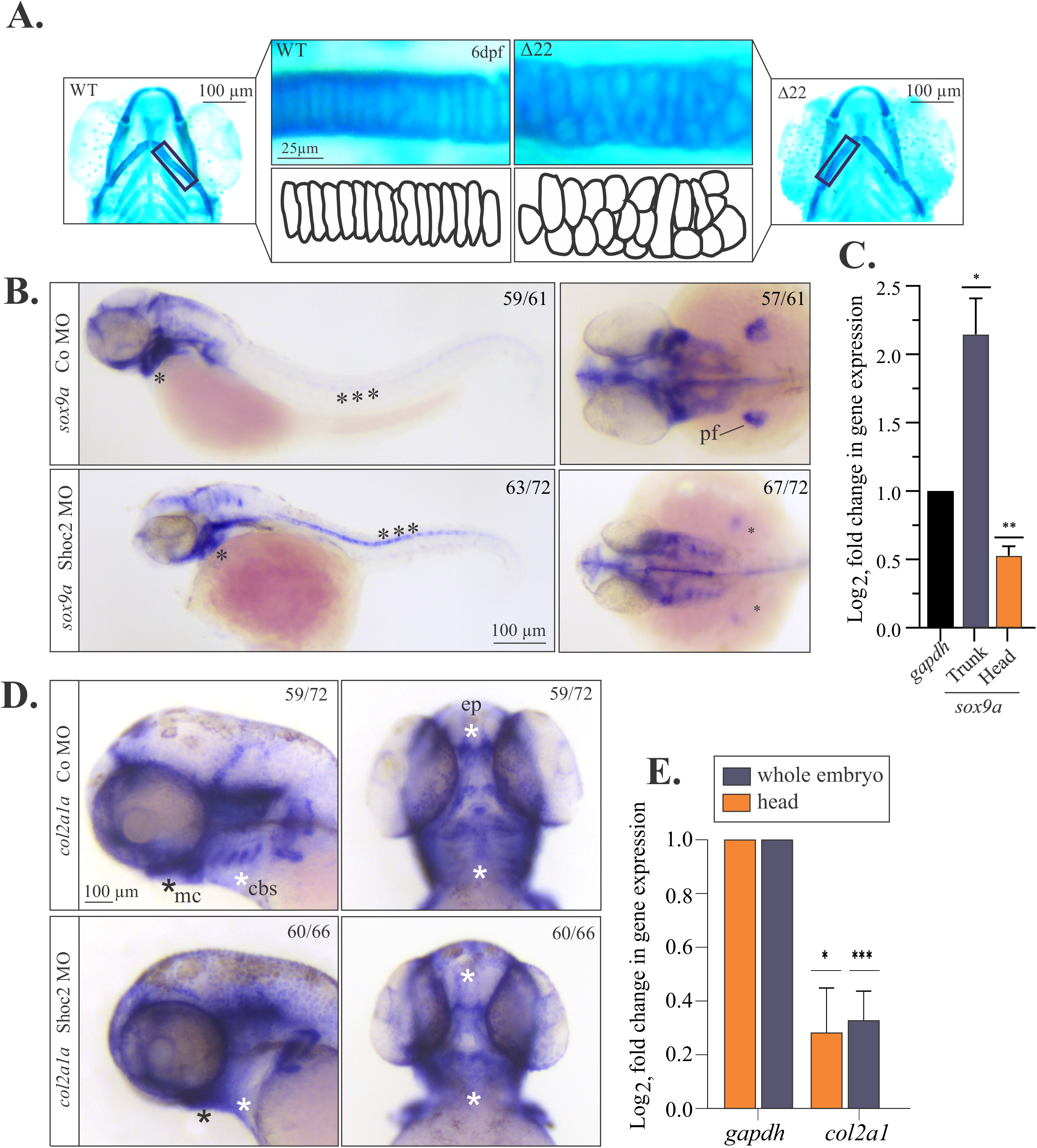
Molecular defects in craniofacial development and chondrocyte morphology in Shoc2 depleted embryos. (**A**) Flat mounts of WT and *shoc2*(Δ22) mutant 6 dpf larvae stained with Alcian blue. The lower panels show individual chondrocytes outlined in black, highlighting the differences in cell morphology of the ceratohyal cartilage. (**B**) Lateral and dorsal views of control and *shoc2* morphant larvae show the *sox9a* expression of 2 dpf larvae. Asterisks show areas of altered *sox9a* expression in the control and *shoc2* morphants. (**C**) Total RNA was extracted from dissected 3 dpf control and *shoc2* morphant larvae. The levels of *sox9a* mRNA expression were quantified by qPCR. *gapdh* is a control mRNA. The data are presented as the Log_2_fold change of the mRNA levels in morphant larvae normalized to control. The results represent an average of three biological replicas. Error bars indicate means with SEM. *p<0.05, ** p<0.01, *** p<0.001 (Student’s t-test). (**D**) Lateral and dorsal views of control and *shoc2* morphant embryos show the *col2a1* expression in 3 dpf larvae. Asterisks mark areas of reduced *col2a1* expression in *shoc2* morphants. (**E**) Total RNA was extracted from dissected 3 dpf control and Shoc2 morphant larvae and levels of *col2a1* mRNA expression were quantified by qPCR. The data are presented as the Log_2_fold change of the mRNA levels in morphant larvae normalized to control. *gapdh* is a control mRNA. The results represent an average of three biological replicas. Error bars indicate means with SEM. *p<0.05, ** p<0.01, *** p<0.001 (Student’s t-test). pf: pectoral fin. mc: Meckel’s cartilage. cbs: ceratobranchials. ep: ethmoid plate.

*Sox9a* directly activates the expression of the alpha I chain of type II collagen (*col2a1*), the major collagen in cartilage and marker of differentiating chondrocytes in zebrafish (Jessen, 2015). When the expression of *col2a1* was compared in control and *shoc2* morphant larvae, dramatic changes in *col2a1* expression were detected in *shoc2* morphants at 3 dpf (**Fig. 5D, E**). Embryos injected with *shoc2* MO exhibited a reduction in *col2a1* staining in the Meckel cartilage, ethmoid plate and ceratobranchial arches (**Fig. 5E**). Other cartilage genes regulated by *sox9a* and required both for proper cartilage formation in the development and maintenance of mature cartilage include proteoglycans aggrecan a and b (*acana and acanb*) (Li et al., 2018, Oh et al., 2014). Similar to *col2a1,* we found dramatic changes in patterns and expression levels of *acana,* and *acanb* in larvae injected with *shoc2* MO (**Fig. 6A, B**). The expression of *acana* and *acanb* was limited to the ethmoid plate from the bilateral cranial NCC streams of the anterior maxillary, Meckel’s cartilage and ceratobranchials elements of *shoc2* morphant larvae (**Fig. 6A, B and Fig. S6A**). qPCR analysis (**Fig. 6C**) further demonstrated that *shoc2* loss leads to the misregulation of *sox9a* signals, thereby affecting the expression of proteins needed for cartilage maturation.

**Figure 6.**
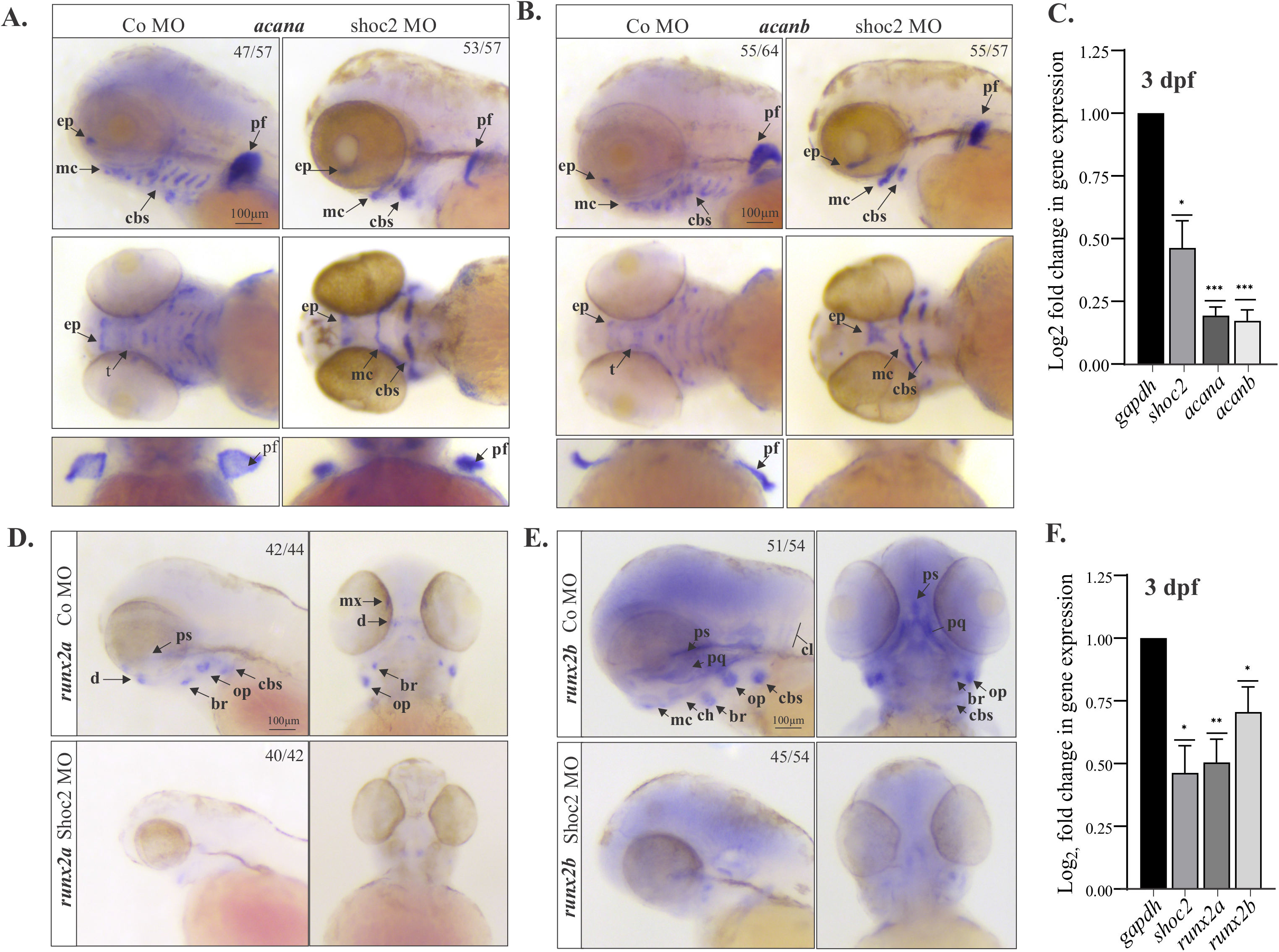
Extracellular matrix and bone pre-cursor genes are deficient in craniofacial cartilage and bone structures in *shoc2* morphants. Lateral and ventral views of control and *shoc2* morphant embryos show expression of the extracellular matrix proteoglycan *acana* (**A**) and *acanb* (**B**) at 3 dpf. Arrows indicate sites of reduced or lost expression. The total number of embryos used in the statistical analysis is indicated on each image. (**C**) Total RNA was extracted from control and Shoc2 morphant larvae at 3 dpf and levels of *shoc2, acana* and *acanb* mRNA expression were quantified by qPCR. *gapdh* is a control mRNA. The data are presented as the Log_2_fold change of the mRNA levels in morphant larvae normalized to control. The results represent an average of three biological replicas. Error bars indicate means with SEM. *p<0.05, ** p<0.01, *** p<0.001 (Student’s t-test). Lateral and ventral views of control and *shoc2* morphant embryos show expression of the *runx2a* (**D**) and *runx2b* (**E**) at 3 dpf Arrows indicate sites of reduced or lost expression. The total number of embryos used in the statistical analysis is indicated on each image. (**F**) Total RNA was extracted from control and Shoc2 morphant larvae at 3 dpf and levels of *shoc2, runx2a* and *runx2b* mRNA expression were quantified by qPCR. *gapdh* is a control mRNA. The results represent an average of three biological replicas. The data are presented as Log_2_fold change of the mRNA levels in morphant larvae normalized to control. Error bars indicate means with SEM. *p<0.05, ** p<0.01, *** p<0.001 (Student’s t-test). ep: ethmoid plate. mc: Meckel’s cartilage. cbs: ceratobranchials. pf: pectoral fin. t: trabeculae. d: dentary. ps: parasphenoid. br: branchiostegal ray. op: opercle. mx: maxilla. pq: palatoquadreate. cl: cleithrum. ch: ceratohyal

To address the extent to which bone ossification is affected in *shoc2* morphants, we assayed the expression of runt-related transcription factors *runx2a* and *runx2b. Runx2a* and *runx2b* regulate the maturation from immature chondrocytes to hypertrophic chondrocytes and osteoblast differentiation during the process of endochondral ossification (Mevel et al., 2019). In embryos injected with control MO, *runx2a* and *runx2b* were expressed in hypertrophic chondrocytes and dermal ossification centers (**Fig. 6D, E**). *Runx2a* was expressed in the cleithrum, dentary, maxilla, operculum, pharyngeal arches and parasphenoid, whereas *runx2b* expression was detected in differentiating osteoblasts of the branchiostegial ray, cleithrum, operculum, palatoquadrate, parasphenoid, and pharyngeal arches (**Fig. 6D, E and Fig. S6A**). Yet, in *shoc2* morphant embryos, *runx2a* and *runx2b* expression was greatly reduced or absent in presumptive cartilaginous elements of the viscerocranium at 3 dpf, indicating that endochondral ossification was practically absent (**Fig. 6F**). These results were consistent with our earlier findings demonstrating a dramatic reduction in calcification of craniofacial bones visualized by Alizarin Red S staining (Jang et al., 2019b) (**Fig. S1A**). Overall, these results demonstrate that Shoc2 function is required for the proper execution of the chondrocyte differentiation program.

### Shoc2 knock-out affects gene expression of the *sox10*-positive cells

Our previous studies established that Shoc2 is expressed in *sox10*-positive cells at 6 dpf (Jang et al., 2019b). Thus, to gain more insight into the transcriptional changes that NC-derived cells experience in the absence of Shoc2, we performed comparative transcriptome analysis of the *sox10*-positive cells. In order to prevent experimental variability associated with MO injections, in these experiments we utilized our CRISPR/Cas9 *shoc2*Δ*22^+/-^* mutant (Jang et al., 2019b) and Tg(*sox10:RFP)* transgenic lines to generate Tg(*sox10:RFP; shoc2*Δ*22^+/-^*) fish. *Sox10*-expressing and *sox10*-derived cell populations were then isolated from 200 embryonic Tg(*sox10:RFP* or Tg(*sox10:RFP; shoc2*Δ*22*) transgenic zebrafish larvae at 6 dpf. Pooled embryos were dissociated and *sox10:RFP^+^* cells were isolated immediately using fluorescence-activated cell sorting (FACS) followed by mRNA isolation and RNA-seq analysis (**Fig. 7A**).

**Figure 7.**
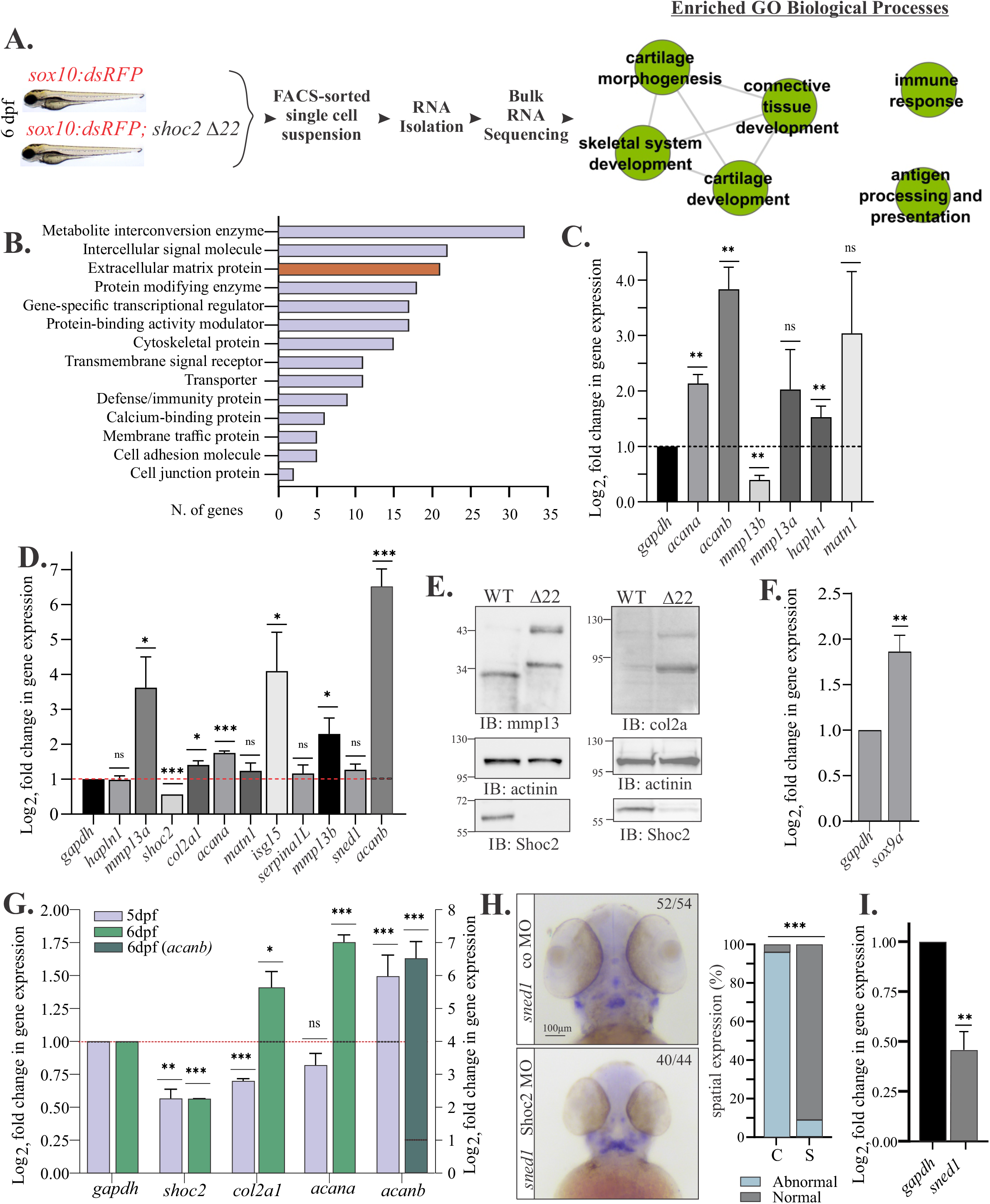
Shoc2 knock-out affects gene expression of the *sox10*-positive cells. (**A**) Cartoon illustrating the workflow used to analyze transcriptome of *sox10*: *RFP* ^+^ cells. Total RNA was isolated from the FACS-sorted *sox10*: *RFP* ^+^ cells, followed by RNA-seq analysis. Data was profiled to identify Enriched GO Biological Processes from Category Compare (Flight et al., 2014). Nodes represent enriched annotation of DEGs. Edges represent relationship between annotation sharing high number of genes with *p*-value cutoff 0.001 and edge weight greater than 0.90. (**B**) The top 15 protein classes of 351 differentially expressed genes (FDR≤ 0.05) analyzed with PANTHER pathway analysis. (**C**) DEGs were selected from the protein class of “Extracellular matrix proteins”. Total RNA was extracted from *sox10*:*dsRed* ^+^ cells of control and *shoc2 null* larvae at 6 dpf and mRNA expression were quantified by qPCR. *gapdh* is a control mRNA. The results represent an average of three biological replicas. The data are presented as the Log_2_fold change of the mRNA levels in the *shoc2 null* larvae normalized to WT larvae. Error bars indicate means with SEM. *p<0.05, ** p<0.01, *** p<0.001 (Student’s t-test). (**D.**) DEGs were selected from the protein class of Extracellular Matrix Proteins. Total RNA extracted from *sox10*:*dsRed* ^+^ control and *shoc2 null* larvae at 6 dpf and levels mRNA expression were quantified by qPCR. The data are presented as the Log_2_fold change of the mRNA levels in the *shoc2 null* larvae normalized to WT larvae. *gapdh* is a control mRNA. The results represent an average of three biological replicas. Error bars indicate means with SEM. *p<0.05, ** p<0.01, *** p<0.001 (Student’s t-test). (**E**) Control and *shoc2 null* larvae were harvested for immunoblotting at 6 dpf. The expression of indicated proteins was analyzed using specific antibodies by WB. (**F.**) Total RNA extracted from WT and *shoc2 null* larvae at 6 dpf. mRNA expression of *sox9a* was quantified by qPCR at 6dpf. The data are presented as the Log_2_fold change of the mRNA levels in the *shoc2 null* larvae normalized to WT larvae. *Gapdh* is a control mRNA. The results represent an average of three biological replicas. Error bars indicate means with SEM. *p<0.05, ** p<0.01 (Student’s t-test). (**G**) Total RNA extracted from control and *shoc2 null* larvae. mRNA expression was quantified by qPCR at 5 or 6 dpf. The data are presented as the Log_2_fold change of the mRNA levels in the *shoc2 null* larvae normalized to WT larvae. *Gapdh* is a control mRNA. The results represent an average of three biological replicas. Error bars indicate means with SEM. *p<0.05, ** p<0.01 (Student’s t-test). (**H**) Ventral view of WT and *shoc2* morphant embryos showing expression of *sned1* in 3 dpf larvae. Aberrant expression patterns of *sned1* are evident in *shoc2* morphants. The graph shows the frequency of abnormal patterns. The total number of embryos used in the statistical analysis is indicated. The results represent an average of three biological replicas. Statistically significant differences between shoc2 MO and control MO according to the Pearson’s chi-squared test are indicated by *p<0.05, **p<0.01, ***p<0.001. (**I**) Total RNA was extracted from control and *shoc2 null* larvae at 6 dpf. The levels of *sned1* mRNA expression were quantified by qPCR at 5 and 6 dpf. The data are presented as the Log_2_fold change of the mRNA levels in the *shoc2* morphant larvae normalized to control larvae. *gapdh* is a control mRNA. The results represent an average of three biological replicas. Error bars indicate means with SEM. *p<0.05, ** p<0.01 (Student’s t-test).

RNA-seq reads were aligned to the *Danio rerio* GRCz11 reference genome (GRCz11.fa) using STAR (version 2.6) (Dobin et al., 2013), followed by the assembly and merging using Cufflinks software package (Trapnell et al., 2012).The number of mapped reads ranged from 27.3 million to 30.3 million per sample and resulted in an overall mapping rate of approximately 97%. Taken together, this indicates both a depth and breadth of sequencing coverage allowing for comprehensive analysis of differentially expressed genes (DEG).

To identify DEGs, data were analyzed using DESeq2 (Love et al., 2014) and CuffDiff (Trapnell et al., 2012). DEGs were then ranked using a false discovery rate (FDR) < 0.05 and fragments per kilobase of exon per million reads mapped (FPKM) ranking resulting in 351 differentially expressed genes, with 188 upregulated and 163 downregulated. The Log_2_fold changes for the obtained gene set are highlighted on the volcano plot (**Fig. S5**).

Further analysis of DEGs for gene ontology biological processes (GO: BP) and KEGG pathways (Kanehisa et al., 2010) identified significant enrichment in biological terms, including “skeletal tissue development”, “cartilage morphogenesis”, “connective tissue development”, and “cartilage development” (**Fig. 7A**). Other significant biological terms included “immune response” and “antigen processing and presentation”. Further analysis of the DEGs by the Protein Analysis Through Evolutionary Relationships (PANTHER) resource separated DEGs into PANTHER protein classes, with “metabolite interconversion enzymes”, “cytoskeletal protein”, “protein modifying enzyme” and “ECM proteins” being the top enriched classes (**Fig. 7B, Table S2**). Together, these analyses suggested that deficiencies in the development of the NC-derived cartilage and bone observed in *shoc2* CRISPR mutants are potentially due to changes in expression of ECM-related proteins. *Sox10*-RFP positive cells from *shoc2* mutant larvae showed robust changes in the expression of ECM proteins (e.g. *acana, acanb*, *sned1, matn1*, *hapln1b*), collagens (e.g. *col6a3*, *col9a1*, *col2a1*, *col11a2/a1, col7a*, *col5a, etc.*), ECM-affiliated proteins (e.g. *anxa1*, *snorc*), ECM regulators (e.g. *mmp13*, *serpinh1*), and other genes associated with ECM remodeling (e.g. *pcolce2, loxl4*) (**Fig. 7B)**. Importantly, many of the DEGs identified in this screen have previously been implicated in being aberrantly expressed in developmental diseases of cartilage or related NC-derived tissues (Carnovali et al., 2019).

Several relatively abundantly expressed genes (i.e. *acana*, *acanb*, *mmp13a, mmp13a*, *hapln1, and matn1)* were selected for additional confirmation by qRT-PCR using mRNA isolated from *sox10:RFP*^+^ cells (**Fig. 7C**). All of the analyzed genes showed consistent expression patterns (Log_2_fold change) between RNA-seq and qRT-PCR analysis. Comparable changes in expression were also observed when the whole larvae were utilized for qRT-PCR analysis (**Fig. 7D**). Interestingly, we found that the expression of ISG15 ubiquitin-like modifier related to the interferon response (Perng and Lenschow, 2018) was significantly upregulated. ISG15 and MMP13 were previously associated with a population of mature-hypertrophic chondrocytes (Wu et al., 2021). Due to the limited availability of antibodies recognizing zebrafish proteins, we examined the protein expression of MMP13a and Collagen 2*a*1 only. Results in **Fig. 7E** confirmed that increased mRNA expression of *mmp13* and *col2a1* corresponded to elevated protein levels of processed and unprocessed forms of MMP13 and Collagen 2*a*1.

Surprisingly, these findings were in contrast to what we observed at 3 dpf, where *shoc2* loss led to a significant decrease in the expression *col2a1*, *acana*, *acanb,* and *sox9a* (**Figs. 5 and 6**). These observations prompted the hypothesis that Shoc2-mediated signals affect the temporal aspect of protein expression during embryonic development. To test this hypothesis, we first established temporal patterns of expression for *col2a1*, *acana* and *acanb* in WT larvae. The qRT-PCR analysis demonstrated that *col2a1, acana,* and *acanb* expression raises significantly around 3 dpf followed by a sharp decrease in RNA expression by 6 dpf, when compared to the expression levels at 2 dpf (**Fig. S6B**). However, when mRNA expression of *col2a1*, *acana* and *acanb* was examined in Shoc2 *null* larvae, we found that the expression levels of *col2a1*, *acana* and *acanb* were increasing gradually, when compared to WT larvae at 5 and 6 dpf (**Fig. 7G**). Likewise, we also found the expression of *sox9a* in Shoc2 *nulls* was much higher than in WT larvae at 6 dpf **(Fig. 7F**). Of note, the expression of *col2a1*, *acana* and *acanb* in Shoc2 *null* larvae at 6 dpf was still much lower than their expression levels detected at 3 dpf of WT larvae (**Fig. S6C**).

An additional DEG evaluated in this study was the structural ECM glycoprotein, SNED1 (Sushi, Nidogen and EGF-like Domains 1). *Sned1* is broadly expressed during development, in particular, in NC and mesoderm derivatives (Vallet et al., 2021), and was previously implicated in multiple aspects of mouse embryonic development, including the formation of craniofacial structures (Barque et al., 2021). Zebrafish SNED1 protein orthologue shares 64% identity with its human counterpart and preserves protein domains found in human SNED1 (Vallet et al., 2021). *sned1* expression was easily detectable in cranial structures of control 3 dpf larvae by WISH, but, was reduced considerably in *shoc2* morphants (**Fig. 7H, I and Fig. S6D**). Of note, our data in **Fig. 7D** show that the expression of *sned1* at 6 dpf was somewhat elevated in *shoc2 null* larvae (**Fig. 7D**). These data indicate that, in the absence of Shoc2, ERK1/2 signals regulating the expression of ECM-related proteins are delayed, further supporting our hypothesis that Shoc2 signals control temporal expression programs during development.

We conclude, that Shoc2-controlled signals regulate the expression of master transcription factors in the gene regulatory network governing the NC developmental program. Aberrantly activated ERK1/2 signaling at the early embryonic stages caused in disbalance extracellular matrix turnover either by decreased matrix synthesis and/or increased matrix degradation which in its turn may drive long lasting defects in organ development (**Fig. 8**).

**Figure 8.**
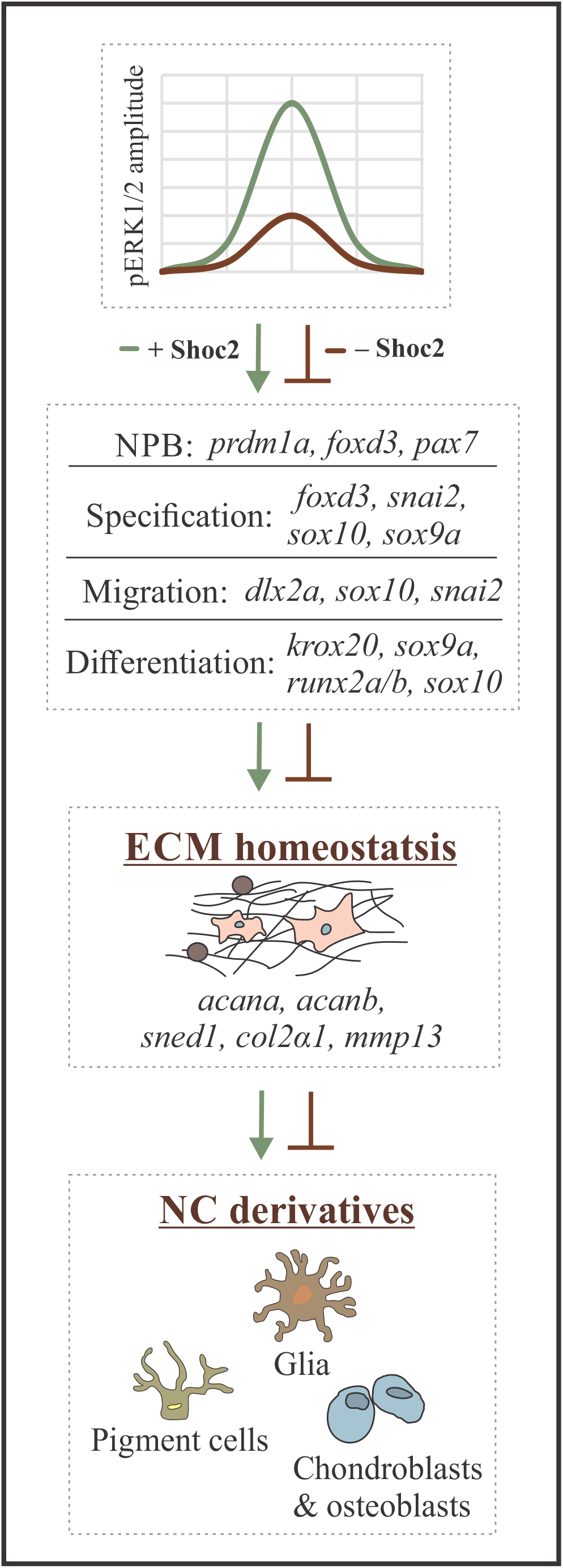
Schematic diagram showing the working model depicting what is currently understood for the role of Shoc2 embryonic development. Shoc2 amplifies and modulates ERK1/2 signaling during early stages of NC development. Lack of Shoc2 followed by the changes in dynamics of ERK1/2 signals (e.g. reduction in signals amplitude) affects expression of key transcription factors essential for NCCs specification, migration and differentiation. Defects in early stages of NC development lead to aberrant expression of downstream effector genes, including proteins regulating homeostasis and remodeling of ECM, ultimately leading to the profound defects in multiple NCC derivatives.

## Discussion

In this study, we have attempted to identify how Shoc2-mediated signals regulate embryogenesis and development of NC for a better understanding of the precise etiology of NSLH and, possibly, other related RASopathies. To analyze how Shoc2 affects the expression of the key regulatory genes participating in the various stages of NC development, we employed MO treatment to block translation from maternal and zygotic *shoc2* mRNAs. MO depletion uncovered developmental deficits obscured by the maternal RNA in the CRISPR/Cas9 Shoc2 *null* mutants (Jang et al., 2019b). We found that loss of Shoc2 affected the expression of the key NPB markers *sox2*, *pax7*, and *prdm1a* at the stage when cells at the NPB commit to the NC fate (**Fig. 1**). Although we have not detected changes in the major-to-minor axis ratio of the *shoc2* morphants, we cannot rule out that Shoc2 signals influence the coordinated convergent extension cell movements during zebrafish gastrulation. Yet, early abnormalities of Shoc2 MO larvae varied from the phenotypes of the zebrafish injected with ERK1 and ERK2 MOs (Krens et al., 2008). Injection of the ERK1 and ERK2 MO severely altered anterior-posterior extension of the dorsal body axis and caused easily detectable deficits in cell movement during gastrulation and low survival rates at 24 hpf (Krens et al., 2008). Future studies will determine whether differences in the observed phenotypes are due to the specificity of the Shoc2-mediated signals or simply changes in the ERK1/2 amplitude.

Loss of Shoc2 also affected the expression patterns of the NC specifier required for the NCCs formation in the NPB, *foxd3* (Lister et al., 2006, Montero-Balaguer et al., 2006, Stewart et al., 2006). The changes in the expression of *foxd3* concurred with the altered expression of other NCC specifiers, *snai2* (Thisse et al., 1995), *sox9a*, and *sox10* (Dutton et al., 2001, Carney et al., 2006, Haldin and LaBonne, 2010), as well as a pan-neural crest marker *crestin* (Rubinstein et al., 2000, Luo et al., 2001) (**Fig. 2**). As was determined by the limited migration of *dlx2a* positive cells and the decreased expression of *sox9a*, *sox10, snai2*, and *crestin* at the 18 ss stage, Shoc2 loss also affects the migration of NCCs (**Fig. 3**). Our data suggest that changes in the expression of the central players regulating the specification of NC (e.g. *foxd3*, *sox9a,* and *sox10)* likely trigger changes in transcriptional programs responsible for cytoskeletal rearrangement, a gain of motility by NCC, and the downstream epithelial-to-mesenchymal transition (EMT) (e.g. expression *twist, chd1, chd2*). These findings are well-aligned with the results of earlier studies demonstrating that signals of the Shoc2-ERK1/2 axis coordinate cell adhesion and movement of cultured cells via controlling the expression of the various proteins regulating these processes (Jeoung et al., 2016, Kaduwal et al., 2015, Young et al., 2013). Others suggested that M-Ras/Shoc2 signals regulate cell motility by coordinating the turnover of E-cadherin at cell-cell junctions and modulating its interaction with p120-catenin (Kota et al., 2019). Although molecular details of how Shoc2-mediated signals regulate NCC motility are yet to be elucidated, our study provides additional evidence for the critical role of Shoc2 in controlling cell migration.

Transcription factors *foxd3*, *sox9a,* and *sox10* also control the ability of NCC precursors to maintain their multipotency, promote survival of NC progenitors and guide the formation of multiple NC lineages (Haldin and LaBonne, 2010, Teng et al., 2008, Nelms and Labosky, 2010). Thus, given the dramatic changes in their expression in *shoc2* morphant larvae, it is not entirely surprising that the loss of Shoc2 hinders the development of NCC derivatives, such as chondrocytes, melanocytes, Schwann cells, and neurons of the peripheral nervous system (**Figs. 4-6**). Importantly, these findings place Shoc2 ahead of *foxd3*, *sox9,* and *sox10* in the gene regulatory network underlying NC development.

The striking changes in the branchial arches, the severe loss of *crestin*-positive cells in the hindbrain at 24 hpf, and the loss of neurons and glia at 3 dpf of *shoc2* morphants could result from the specific loss of hindbrain premigratory NCCs by cell death. Shoc2 was reported to regulate the survival of neural progenitor embryonic stem cells and the maintenance of their self-renewal capacity (Moon et al., 2011). Although we do not know the specific downstream effectors of *shoc2*-mediated cell death, our data suggest that members of the Snail family of transcriptional repressors are possible candidates. Snail proteins act as survival factors in progenitor cell populations and are often overexpressed in human cancers (Barrallo-Gimeno and Nieto, 2005, Buyuk et al., 2022, Wu et al., 2005, Inoue et al., 2002). A detailed understanding of the role of the Shoc2-ERK1/2 axis in cell death warrants additional studies. The broad range of affected NC lineages and the early reduction in the expression of NC specification genes suggest that Shoc2 might be required to maintain an undifferentiated pool of NC progenitors. Thus, it is tempting to speculate that when Shoc2 is depleted, and only residual ERK1/2 signals regulate NC induction and specification, the differentiation of progenitor cell precursors stall because fewer cells of each type are formed.

One of the most unexpected discoveries is the results of the comparative transcriptome analysis of *sox10*-positive cells and our findings that the persistent changes in the expression of genes coding the various proteins associated with ECM (**Fig. 7**) could be a direct cause of the defects/delays in the ossification of cartilage. Shoc2 loss triggered the deregulation in the expression of many relevant chondrogenesis genes, including *col2a1*, *acana*, *acanb*, *hapln* and *mmp13*, *matn1, and sox9*. We found that the build-up in ECM proteins in *shoc2 nulls* is accompanied by the abnormal expression of ECM regulators like MMP13 and modifying enzymes like *pcolce2* and *loxl4* (**Table S2**). Thus, it is plausible that an accumulation of ECM collagens and glycoproteins hinders the chondrocyte potential to convert into osteoblasts. Concomitant with alterations in the intracellular signaling, metabolite interconversion, and the expression of enzymes modifying ECM proteins (**Fig. 7B and Table S2**), these data point out a global effect of Shoc2-ERK1/2 signals on ECM homeostasis and/or remodeling. Importantly, our data also suggest that during embryonic development Shoc2 scaffold modulates the parameters of ERK1/2 signaling dynamics. Future studies using different methodologies are needed to address whether marked increases in the expression of Col2a1 and aggrecan a and b observed in 5 and 6 dpf *shoc2 null* larvae result from changes in ERK1/2 amplitude or duration. Considering the diverse role collagens and other ECM-related proteins play in NC migration and morphogenesis (Jessen, 2015), it is tempting to speculate that other abnormalities of Shoc2 *null* zebrafish (e.g. vasculature, hematopoiesis) or even abnormalities of NSLH patients are related to the deficits in the homeostasis of the ECM.

Shoc2 has been suggested as an independent prognostic marker and a target for sensitization of the MEK inhibitors in cancer (Jones et al., 2019, Sulahian et al., 2019). Thus, our findings that Shoc2-ERK1/2 signals control ECM homeostasis/remodeling have potential implications for understanding the role of the Shoc2 in tumor progression. In cancer, abnormal ECM dynamics occur due to disrupted balance between ECM synthesis and secretion and altered expression of matrix-remodeling enzymes (Henke et al., 2019). Thus, our findings are directly relevant to the efforts toward novel direct therapeutics targeting the Shoc2-ERK1/2 axis.

In summary, we have characterized a novel and critical role for Shoc2 during NC development. Significant defects in several NC-derived tissues of *shoc2* mutants and morphant larvae indicate that Shoc2 is an essential component of the gene regulatory network guiding the development of NC-derived tissues. Further experiments that will resolve other aspects of the tissue-specific role of *shoc2* during development and in the etiology and pathogenesis of NSLH are in progress to generate a comprehensive model of Shoc2 function.

## ACKNOWLEDGMENTS

We thank Drs. Tianyan Gao, Kristin Artinger, Louis Hersh, Ann Morris and Charles Waechter for critical reading of the manuscript; Kristin Artinger for providing critical reagents and members of Galperin lab for productive discussion.

## AUTHOR CONTRIBUTIONS

E.G. conceptualized and supervised the project; K.A. and E.C.R. analyzed all the RNA sequencing data, L.A. performed the microinjection and western blot experiments. R.G.N. performed in situ experiments and analysis, RT-qPCR experiments and analyses; D.L. performed PH3 immunostaining and in situ experiments, O.T. performed in situ analysis; R.G.N. and E.G. wrote the manuscript with inputs from all authors.

## MATERIALS AND METHODS

### Resource Availability

#### Lead contact

Further information and requests for resources and reagents should be directed to and will be fulfilled by the Lead Contact, Emilia Galperin.

### Materials Availability

All antibodies, chemicals and zebrafish lines used in this study are commercially available. All other unique materials are available upon request.

### Data and code availability

Accession IDs provided by GEO: GSE198231

### Zebrafish strains and maintenance

All zebrafish (*Danio rerio*) strains were bred, raised, and maintained in accordance with established animal care protocols for zebrafish husbandry. Embryos were staged as previously described (Kimmel et al., 1995). All animal procedures were carried out in accordance with guidelines established by the University of Kentucky Institutional Animal Care and Use Committee. Briefly, zebrafish embryos were raised at 28.5°C and kept in 14/10h light/dark cycle and staged according to Kimmel *et al*., 1995 (Kimmel et al., 1995). When necessary, 1-Phenyl-2-thiourea (0.002%) was added to the embryo media to prevent pigment development. *Shoc2*Δ*22* zebrafish were maintained as heterozygotes and incrossed to generate homozygous mutant embryos. The *Shoc2*Δ*22* heterozygous mutant line was crossed with available reporter lines Tg(*sox10*:*RFP*). The mutant *shoc2* zebrafish line Shoc2 E2Δ22+/-(ZDB-GENE-050208-523) was reported previously (Jang et al., 2019b).

### Morpholino and mRNA injection

All MOs were obtained from Gene Tools, LLC (Philomath, OR) and injected into 1-2 cell stage zebrafish embryos. The following MOs were used in this study: standard control MO: 5’-CCTCTTACCTCAGTTACAATTTATA-3’; *shoc2* MO1: 5’-TACTGCTCATGGCGAAAGCCCCGCA-3’. Embryos were injected with 5.2 ng each of MOs.

### Genotyping

Genomic DNA was extracted from individual embryos or adult tail clips. Briefly, 20 µl of the ThermoPol Buffer (New England Biolabs, # B9004S) was added to the samples and boiled for 5 min. Samples were digested with 50µg (5µL) Proteinase K (Millipore Sigma, #p22308) for 12 hours at 55 °C. Proteinase K was then inactivated by boiling for 10 min. PCR was carried out in a 25 µl reaction solution containing: 1 µl of 10 mM dNTP, 1 µl of 10 mM forward and reverse primer, 2.5 µl of 1x ThermoPol buffer and 0.5 units of Taq Polymerase (New England Biolabs, #M0267). The s*hoc2*Δ*22* heterozygous mutant allele was detected using the primers forward 5’-CCATCAAGGAGCTGACCCAG-3’and reverse 5’-AGTCAGGTAGGCTGGTCAGA −3’.

### Phenotype analysis

#### Skeletal stain

For Alcian blue staining, zebrafish larvae were fixed in 4% paraformaldehyde for 2 h at room temperature and stained according to Kimmel et al., 1998. Calcified structures were examined by acid-free Alizarin Red S staining. Larvae were fixed in 4% PFA for 2 h and stained in a 0.05% Alizarin Red S solution for 30 min in the dark on low agitation. Larvae were then rinsed in a 50% glycerol, 0.1% KOH solution to remove excessive staining and kept at 4◦C in the same solution for imaging.

#### In situ hybridization

Linearized plasmid DNA was cleaned using DNA Clean & concentrator-5 (Zymo Research #D4014). *In situ* hybridization was completed according to a standard protocol (Cunningham and Monk, 2018) using DIG RNA labeling kit T7/SP6 (Millipore Sigma #11175025910). RNA probes were cleaned using SigmaSpin Sequencing Reaction Clean-UP (Millipore Sigma #S5059-70EA). Briefly, embryos were permeabilized with Proteinase K and hybridized in probes diluted to 3 ng/μl overnight at 65-68°C. The probe was removed and embryos were washed and blocked for a minimum of 1 hour prior to incubation in anti-digoxigenin-AP, Fab fragments 1:2,000 (Roche, #11093274910). Signal was detected using NBT:BCIP (1.38:1.0 ratio) (Roche, NBT #11383213001 & BCIP #11383221001). Background staining was removed using brief five-minute washes with methanol. Embryos were cleared through a glycerol series and imaged on the system listed below.

Regions of *sned1* were amplified from cDNA using the primers forward 5’-CATTACTCCCAGGTCAGATGTAC-3’ and reverse 5’-TCAGGCTTAATGCGGTGTCT −3’*. This* amplified region was cloned into the pJET1.2/blunt Cloning Vector (ThermoFisher, #K1232) to generate both sense and anti-sense probes. Additional antisense DIG conjugated probes were synthesized from plasmids kindly gifted from: *collagen2a1* (Dr. Tatjana Piotrowski, The Graduate School of the Stowers Institute for Medical Research, Kansas City, MO), *acana* & *acanb* (Dr. Adele Faucherre, Institut de Génomique Fonctionnelle, Montpellier, France), *sox10* (Dr. Rebecca Cunningham, Washington University in St. Louis, St. Louis, MO), and c*restin, dlx2a, foxd3, krox20, mbp, pax7, prdm1a, snai2, sox2, sox9a, runx2a,* and *runx2b* (Dr. Kristin Artinger, University of Colorado Anschutz Medical Campus, Aurora, Colorado).

#### TUNEL

Apoptotic cells in whole body embryos were detected using Millipore Sigma ApopTag Red *In Situ* Apoptosis Detection Kit S7165. TUNEL was performed as described in (Parada-Kusz et al., 2018). Briefly, embryos were fixed in 4% PFA/PBS, washed through a PBS/methanol gradient ending in 100% methanol and incubated at −20°C for at least one hour. Embryos were then sent through a PBSTw (0.1% tween-20 in PBS) gradient wash ending in 5 x five-minute washes in PBSTw. Embryos, 24 and 48 hpf were permeabilized for 4 or 27 minutes respectively in 20µg/100µL proteinase K (GoldBioP-480-1) prior to being re-fixed in 4% PFA/PBS. Whole embryos were incubated in 50uL equilibrium buffer for 15 minutes at 37°C before the 4 °C overnight incubation in the reaction mix (20uL equilibrium buffer, 12uL reaction buffer, 6uL TDT, 10% Tween-20). Five washes with PBST were completed before adding the stop/wash buffer for 5 minutes. Embryos were then blocked for 1 hour. Finally, anti-DIG solution in blocking buffer was added to embryos and incubated for 30 minutes in the dark. Anti-DIG was removed and embryos were washed PBST and incubated for 1 hour at 37°C in the Fluorescein in the dark. Images were captured as described below.

#### Iridophores

Incident light images of four-day old embryos’ iridophores were captured with a Leica M165FC microscope. All iridophores in a 1350 um long region in the tail (spanning approximately 11 somites) were quantified.

#### Phospho-Histone H3 Immunolabeling

Briefly, embryos were fixed in 4% PFA/PBS for 1 hour at room temperature. Embryos were washed in water for 5 minutes and then incubated one hour at room temperature in a blocking solution (2% goat serum, 1% BSA, 1% DMSO, 0.1% Triton-X-100, 1X PBS). Embryos were then incubated overnight at 4°C in primary antibody diluted in a blocking solution (Anti-phospho-Histone H3 (Ser10) Antibody, Mitosis Marker, Millipore Sigma, 06-570). Embryos were thoroughly rinsed in PBS-Triton-X (0.1%). Next, embryos were incubated in the dark overnight in an Alexa488 conjugated secondary antibody diluted in blocking solution (1:750). Finally, embryos were rinsed with 0.1% Trition-X-100 and imaged.

#### Imaging methods and analysis

Images of whole-mount *in situ* hybridization and whole body Alcian blue staining were captured with a Leica DFC450 digital camera. Alcian blue ceratohyal images were acquired with a Zeiss Imager AzioCam MRm. Fluorescent images from methods TUNEL and pH3 immunolabeling were captured with a Leica M165FC microscope.

### Molecular analysis

#### Real-time quantitative polymerase chain reaction (RT-qPCR)

Total RNA was isolated from the pool of 25 embryos (or dissected embryo tissue) using PureZOL RNA Isolation Reagent (Bio-Rad, #732-6890) and Aurum Total RNA Isolation Kit (Bio-Rad, #732-6820). Aliquots containing equal amounts of RNA were subjected to RT-PCR analysis (Bio-Rad, iScript™ Reverse Transcription Supermix for RT-qPCR, # 1708840). qPCR was performed using Bio-Rad iTaq™ Universal SYBR® Green Supermix (#1725120) and a Bio-Rad CFX detection system (Bio-Rad, CA). Relative amounts of RNAs were calculated using the comparative C_T_ method. Sequence-specific primer sets are presented in **Table S1**. The values for the samples were normalized against those for the reference gene, and the results are presented as the Log_2_fold change in the amount of mRNA recovered from WT and mutant embryo. The data represent the means ± SEM from three independent experiments.

#### Western blot analysis

Proteins were extracted from de-chorionated and de-yolked embryos/larvae and resolved by SDS-PAGE. In short, water was removed from approximately 25 embryos in a microcentrifuge tube. One solid glass bead and 50 µL RIPA buffer containing protease inhibitors were added to the embryos. Gentle manual agitation physically lysed the embryos. Samples were centrifuged at 4 °C for 10 minutes at 14,000 RPM. Total protein lysate was removed from pellet and bead. 25µg of total lysate per sample was run on a 10% acrylamide gel. Western blot analysis was performed as described previously (Wilson et al., 2021). Quantification was performed using the densitometry analysis mode of Image Lab software (Bio-Rad, CA). Antibodies against the following proteins were used: MMP13 polyclonal antibody: Proteintech. #18165-1-AP. Collagen Type II. Developmental Studies Hybridoma Bank. #II-II6B3. Anti-α-actinin Antibody (H-2). Santa Cruz. #17829. Anti-Sur-8 Antibody (E-4). Santa Cruz. #514886. Anti-β-Actin Antibody (C4). Santa Cruz #47778.

#### RNA-seq analysis

Zebrafish transgenic larvae were homogenized and fluorescence-activated cell sorted. Briefly, 6dpf embryos were dissociated in trypsin using a 20G needle and incubated for 2 minutes at 37°C. This was repeated four times. Dissociated cells were strained through a 50µm strainer into 2mM EDTA/5%goat serum/PBS and centrifuged for 10 minutes at 3,500rpm. The cell pellet was resuspended in 1mM EDTA/10% goat serum/PBS. Cells were sorted for RFP+ identity at the University of Kentucky Flow Cytometry and Immune Monitoring Core at the Markey Cancer Center. After sorting, cells were centrifuged at 3,000rmp for 10 minutes. Supernatant was discarded and the RFP+ cells were frozen (−80°C) in PureZol (Bio-Rad. #732-6890). Triplicates of RNA from RFP+ cells were purified as described above.

For library preparation, mRNA was first extracted from total RNA using oligo (dT) magnetic beads and sheared into short fragments of about 200 bases. The cDNA library was sequenced using Illumina NextSeq 500 sequencer. Quality control (QC) of the raw sequence data was performed using FastQC (version 0.11.7). The concatenated sequences were directly aligned to the Danio rerio GRCz11 reference genome assembly (GRCz11.fa) using STAR (version 2.6), generating alignment files in bam format. The alignment rate for each sample is above 90%. Fragments per kilobase per million mapped (FPKM) reads were determined for all RefSeq genes using CuffDiff 2 (FDR ≤ 0.05). For the Cuffdiff2 analysis, Cuffnorm was used to produce FPKM (Fragments Per Kilobase Million) normalized counts. The counts were then filtered to include only genes with minimum expression of one FPKM in three or more samples and an average expression of at least one FPKM. The RNA-seq data is publicly available as GEO series GSE198231. The data is MIAME compliant (https://www.ncbi.nlm.nih.gov/geo/query/acc.cgi?acc=GSE198231).

#### Gene ontology (GO) and pathway and network analysis

Differentially expressed genes determined by RNA-seq analysis were used for functional enrichment including the Category Compare that predicts molecular and cellular functions using the Ingenuity Knowledge base as the background. Gene ontology terms within the data set were provided by Protein Analysis Through Evolutionary Relationships (Thomas et al., 2003).

